## Supplementary material for "Wikipedia as a tool for contemporary history of science: A case study on CRISPR": Benjakob et al., SupFigures

### Code accessibility

Our code for the corpus builder can be found at:
https://github.com/RonaTheBrave/WikiCorpusBuilder

The article’s data is accessible at https://zenodo.org/record/7206381#.Y1JoEezP23I
DOI:10.5281/zenodo.7206381

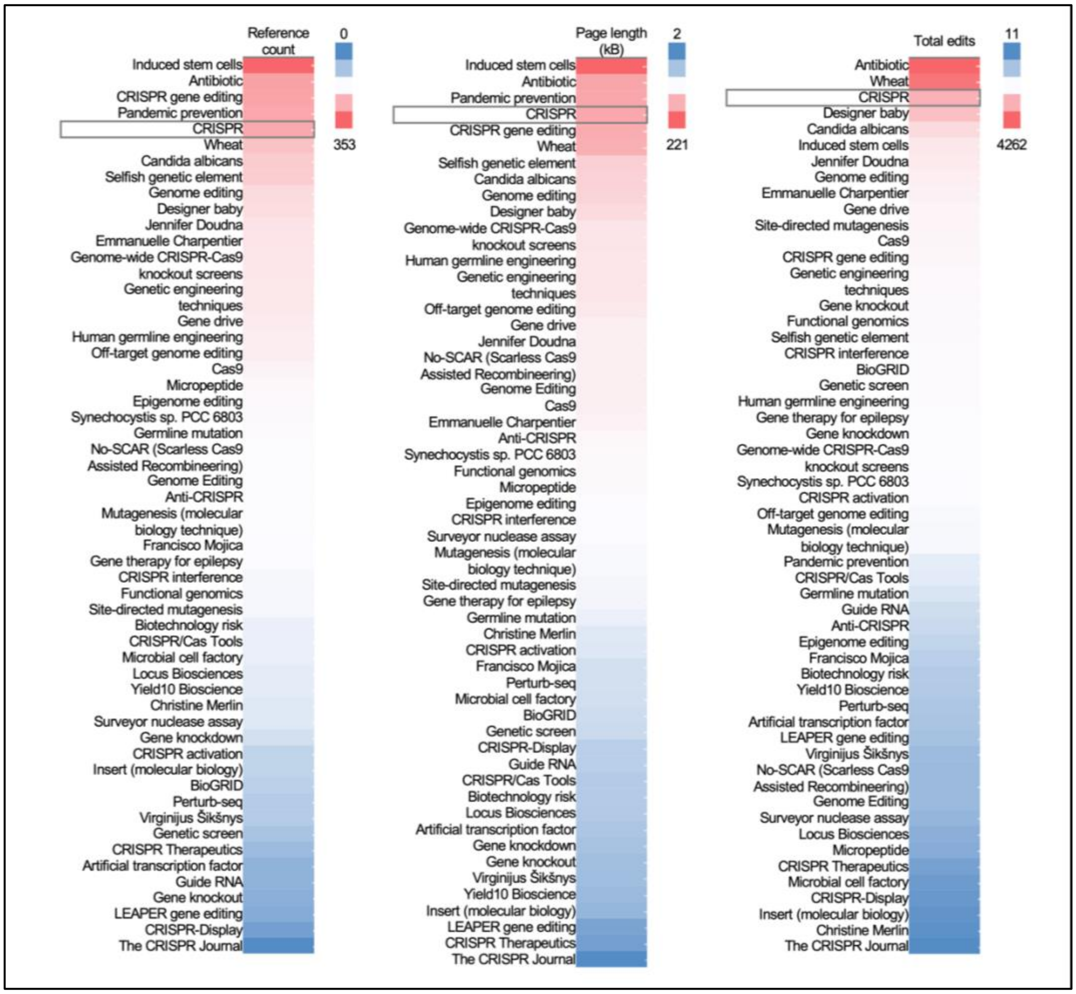


**Fig S1. The CRISPR corpus in numbers.** The articles included in the corpus, sorted by number of references, size in kilobytes (kB) and number of edits. “CRISPR”, highlighted, was among the top 5 articles of each category.

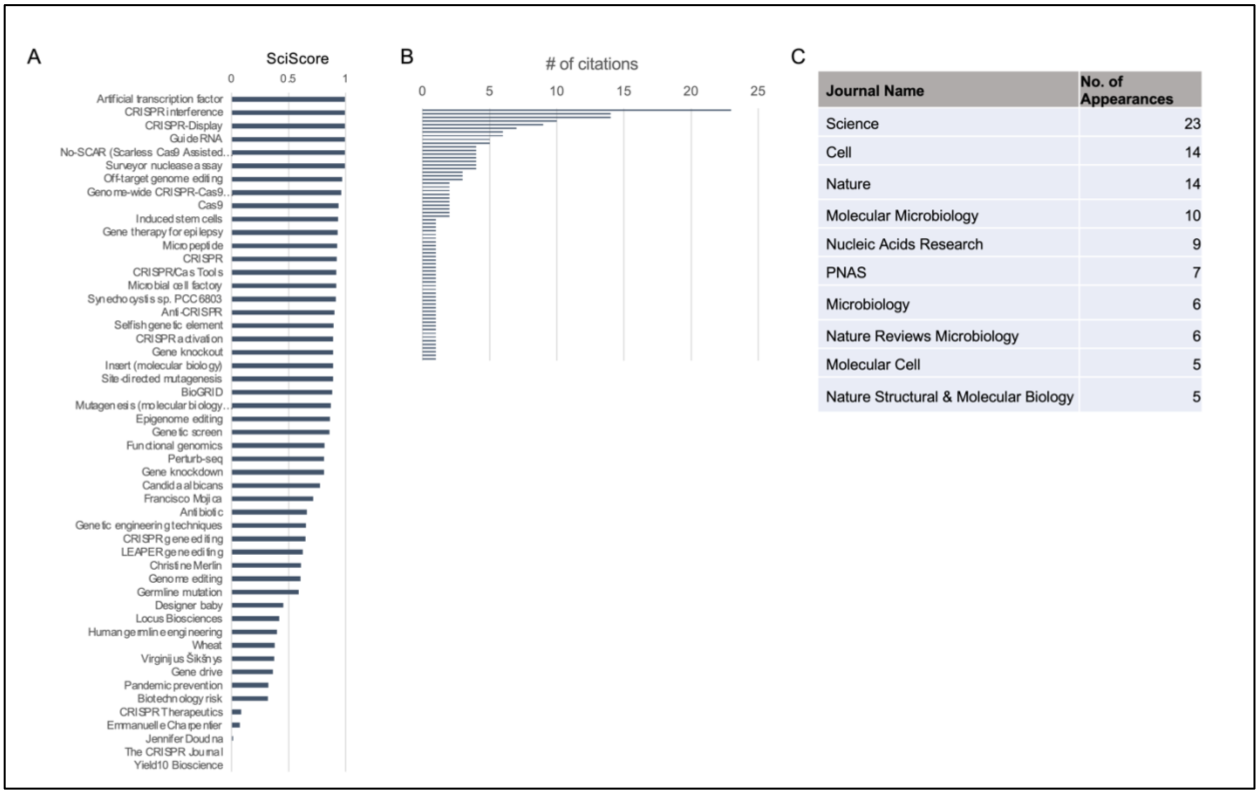


**Fig S2. CRISPR article's references.** A) The corpus SciScore. B) Peer-reviewed journals cited as references in the article as of June 2022, sorted by the number of references per publication. C) A list of the top cited journals (from B) with ≥5 appearances.

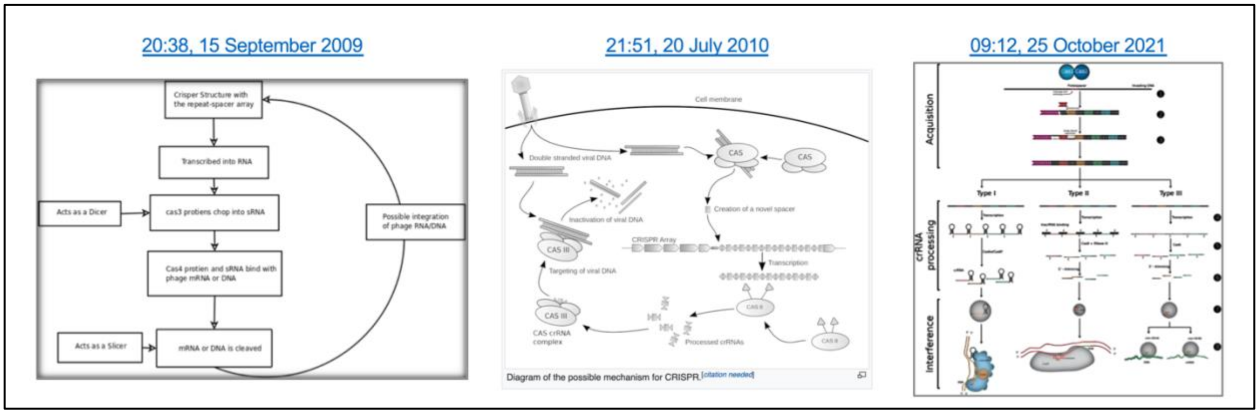


**Fig S3. Illustrations of the CRISPR model.** Shown are a selection of screen grabs from the CRISPR article, reflecting the evolution of Wikicommons graphics of CRISPR's mechanism of action and key players. These are of different versions of the same illustration (A and B) and of a third illustration added later to the article.
